# Age-related phenotypes in breast cancer: a population-based study

**DOI:** 10.1101/2023.05.22.541427

**Authors:** Amalie A. Svanøe, Rasmus O.C. Humlevik, Gøril Knutsvik, Anna K.M. Sæle, Cecilie Askeland, Lise M. Ingebriktsen, Ulrikke Hugaas, Amalie B. Kvamme, Amalie F. Tegnander, Kristi Krüger, Benedicte Davidsen, Erling A. Hoivik, Turid Aas, Ingunn M. Stefansson, Lars A. Akslen, Elisabeth Wik

## Abstract

Breast cancer in young (<40 years) is associated with a higher frequency of aggressive tumor types and poor prognosis. It remains unclear if there is an underlying age-related biology that contributes to the unfavorable outcome. We aim to investigate the relationship between age and breast cancer biology, with emphasis on proliferation. Clinico-pathologic information, immunohistochemical markers, and follow-up data were obtained for all patients aged <50 (Bergen cohort-1; n=355) and compared to previously obtained information on patients aged 50-69 (Bergen cohort-2; n=540), who participated in the Norwegian Breast Cancer Screening Program. Data from the Molecular Taxonomy of Breast Cancer International Consortium (METABRIC) was applied for validation and for analyses of gene expression signatures. Young breast cancer patients presented more aggressive tumor features such as hormone receptor negativity, HER2 positivity, lymph-node metastasis, the HER2-enriched and triple negative subtypes, and shorter survival. Age <40 was significantly associated with features of stemness and higher proliferation (by Ki67), also in molecular subsets. Ki67 showed weaker prognostic value in young patients. We point to aggressive phenotypes, increased tumor cell proliferation and stem-like features in breast cancer of the young. Hence, tumors of young breast cancer patients may have a unique biology, also when accounting for screen/interval differences, that may open for new clinical opportunities, stratifying treatment by age.

## Introduction

Breast cancer incidence rises with age^1^ ^2^, but 7 % of breast cancer patients in high-income countries are diagnosed at age ≤40. Breast cancer of the young, defined as age below 40 or 45 years at diagnosis, frames a cluster of patients with specific age-related challenges (e.g. long-term sequelae after therapy, fertility preservation, genetic counselling)^3^. Among women in this age group, breast cancer is the most common cancer type, and is one of the most frequent causes of death^4^ ^5^.

Young breast cancer patients have poorer prognosis than older patients^6^ ^7^ and higher frequencies of aggressive clinico-pathologic features^8^ ^9^. Previous studies have demonstrated molecular alterations in age-specific patterns^10^ ^11^. Inherited breast cancer based on germline *BRCA1* and *BRCA2* mutations affects the younger patients more frequently. After adjusting for the BRCAness and molecular subtypes, additional age-related biologic alterations are suggested, underpinning the increased cancer aggressiveness seen in breast cancer of the young^12^. Also, a higher tumor proliferation is observed in the younger patients^7^ ^11^ ^13^.

Breast cancer of the young is understudied with respect to age-related biomarker- and context-dependent biologic differences. More studies are needed to evaluate whether there is a need to adjust for age-related differences when tailoring therapy and follow-up protocols for breast cancer of the young. By studying population-based in-house cohorts and external gene expression data, we explored age-related breast cancer phenotypes and clinical outcome, also comparing breast cancer of the young with patients diagnosed in a breast cancer screening program. Our study indicates a weaker prognostic value of Ki67 in the young and supports also other age-related biological breast cancer alterations with clinical relevance. We propose increased clinical attention to patients diagnosed with breast cancer at age below 40 years.

## Materials and methods

### Patient in-house cohorts

A breast cancer patient series (Bergen cohort-1) was established, including tumor tissue and clinico-pathologic data from all women below the age of 50 years, residing in Hordaland County, Norway, diagnosed with primary invasive breast cancer during the period 01.01.1996-31.12.2003. Hordaland County, with 500 000 inhabitants, represents about 10 % of the total Norwegian population. Information on the identity of these patients was obtained through the local pathology registry (Dept. of Pathology, Haukeland University Hospital) and the Cancer Registry of Norway^14^. A previously thoroughly described series of breast cancer patients aged 50-69 years at time of diagnosis was used for comparative analyses (n=543, Bergen cohort-2)^15^ ^16^. This patient series consisted of a population-representative selection of women diagnosed with primary invasive breast cancer in Hordaland County, who participated in the prospective Norwegian Breast Cancer Screening Program (NBCSP) during the same period. In the Bergen cohort-2, 74.2% (403/543) of the carcinomas were detected by screening while the remaining 25.8% (140/543) were detected in an interval between two screenings. Patients with distant metastatic disease at the time or within six months of diagnosis (stage IV) were not included.

#### Bergen cohort-1

In the Bergen cohort-1, 378 breast cancer cases below 50 years of age at time of diagnosis were identified in the specified period. For various reasons, 23 patients were excluded **(Supplementary Figure 1**). This left 355 cases for further analyses. Of these, eight presented with metastasis at time of diagnosis or within six months after the primary breast cancer diagnosis. These were excluded from analyses when comparing the Bergen cohort-1 and -2, and in the survival analyses. The patients received treatment according to standard national protocols for the period of diagnosis.

#### Clinico-pathologic variables, Bergen cohort-1

Clinico-pathologic variables were obtained from clinical and histopathologic records and included patient age at diagnosis, tumor diameter, histologic type and grade, lymph node status, and hormone receptor status. Histologic type was assessed according to WHO criteria, whereas histologic grade was evaluated using the Nottingham grading system^17^. Tumor size was primarily assessed by histological methods (83.6%). When pathologic tumor size was not available (as in patients with locally advanced or multifocal disease), the radiologic or clinical size estimate was included (4.7% and 11.7 %).

#### Specimen characteristics, Bergen cohort-1

For the 355 cases in the Bergen cohort-1, tumor tissue blocks were available for 339 of the cases (95.5%); obtained from surgery (n=288, 85.0%), open biopsy (n=33, 9.7%), or core needle biopsy (n=18, 5.3%) when surgical specimens were not available. The tissue was fixed following standard protocol at the time, using 4% buffered formaldehyde before processed and further paraffin embedded. Storage of the formalin fixed paraffin embedded (FFPE) blocks were up to 27 years (20-27 years).

#### Tissue microarray (TMA), Bergen cohort-1

Tissue microarrays (TMAs) were made in cases with sufficient tumor tissue available (n=319), using a semi-automated instrument (Minicore 3, Tissue Arrayer, Alphelys, France). Haematoxylin and eosin stained slides were evaluated to find areas of highest histologic grade and the peripheral invasive front. Three one mm punches from the corresponding tumor area in the FFPE tissue block were mounted in a recipient paraffin block. From the TMA-blocks, 4–5 μm sections were cut and mounted on poly-lysine coated glass slides. Whole sections (WS) were added for cases where sufficient tumor tissue was not available.

#### Follow-up data

The follow-up information for both cohorts was acquired from the Norwegian Cause of Death Registry and can be considered accurate and complete, and included survival status, survival time and cause of death. The last follow-up date of was 30.06.2017, with a median follow-up time of survivors of 209 months (range 162-257 months), for both cohorts. During the follow-up period, 182 patients (20.5%) died from breast carcinoma and 133 (14.9%) died from other causes.

#### Ethical approval

The study was approved by the Western Regional Committee for Medical and Health Research Ethics, REC West (2014/1984/REK vest). Written informed consent was waived by the ethics committee. The national identification numbers of all patients were checked with the Registry of Withdrawal from Biological Research Consent by the Norwegian Institute of Public Health. None of the cases were listed in the registry. For the Bergen cohort-2, all participants were additionally contacted with written information on the study and asked to respond if they wanted to be excluded. In total, 9 patients (1.6%) withdrew their consent and were excluded from the study. The study was performed in accordance with the declaration of Helsinki.

### Immunohistochemistry

To classify molecular subtypes in the Bergen cohort-1, Estrogen receptor (ER), Progesterone receptor (PR), HER2, Ki67, and Cytokeratin 5/6 (CK5/6) were stained by immunohistochemistry (IHC) and evaluated. When TMA was not available (se section “TMA Bergen cohort-1”), whole sections were used for the remaining cases with invasive tumor for complete assessment of HER2 (n=13), CK5/6 (n=17) and Ki67 (n=12). ER and PR status was obtained from pathology reports in 330 and 329 cases and by new IHC in 24 and 25 cases. For cases in the Bergen cohort-2, the subtypes were classified by the same method^15^ ^16^.

#### Immunohistochemistry protocols

IHC was performed on 4–5 μm sections from FFPE TMA-blocks and on whole section slides in cases with insufficient material for TMA. All slides were de-waxed in ethanol and xylene.

##### Ki67

Staining procedures were performed with a DAKO autostainer using the K4061/Envision Dual Link System (rabbit + mouse). Target retrieval was done in a pressure cooker (Decloaking Chamber Plus, Biocare Medical) in Target Retrieval Solution pH 9.0 (S2367, DAKO). The slides were pre-treated by incubating 8 minutes with Peroxidase Blocking Reagent (S2001, DAKO). A monoclonal rabbit Ki67 antibody (M7240, clone MIB-1, DAKO) was incubated for 30 minutes at room temperature at a 1∶100 dilution, followed by Labelled Polymer-HRP Anti-Mouse for 30 minutes. Finally, diaminobenzidine (DAB) as chromogen was added for 10 minutes and haematoxylin (S3301, DAKO) as a counterstain for 10 minutes. Tissue slides were stored for shorter than two weeks at 4 °C until Ki67 staining was performed.

##### ER and PR

Staining procedures were performed with a DAKO autostainer, using the K4061/Envision Dual Link System (rabbit + mouse). Target retrieval was done by boiling the slides in Target Retrieval Solution pH 9.0 (S2367, DAKO) in a microwave with 6th Sense function (Whirlpool). The slides were pre-treated by incubating 8 minutes with Peroxidase Blocking Reagent (S2001, DAKO). For ER, the monoclonal mouse antibody (M7047, Clone 1D5, DAKO) was incubated for 30 minutes at room temperature at a 1∶50 dilution. For PR, the PgR monoclonal mouse antibody (M3569, clone 636, DAKO) was incubated 30 minutes at room temperature at a 1:150 dilution. Both were followed by Labelled Polymer-HRP Anti-Mouse for 30 minutes. Finally, diaminobenzidine (DAB) as chromogen was added for 10 minutes and haematoxylin (S3301, DAKO) as a counterstain for 10 minutes.

##### CK5/6

Staining procedures were performed with a DAKO autostainer using the K4061/Envision Dual Link System (rabbit + mouse). Target retrieval was done in a pressure cooker (Decloaking Chamber Plus, Biocare Medical) in Target Retrieval Solution pH 9.0 (S2367, DAKO). The slides were pre-treated by incubating 8 minutes with Peroxidase Blocking Reagent (S2001, DAKO). For CK5/6, the monoclonal mouse antibody (M7237, Clone D5/16 B4, Dako) was incubated for 30 minutes at room temperature at a 1∶200 dilution followed by Labelled Polymer-HRP Anti-Mouse for 30 minutes. Finally, diaminobenzidine (DAB) as chromogen was added for 10 minutes and haematoxylin (S3301, DAKO) as a counterstain for 10 minutes.

#### Evaluation of staining

##### Hormone receptors

Tumors were classified ER and PR positive when ≥ 10% of tumor nuclei stained positive, according to guidelines of the period of diagnosis and treatment (1996-2003).

##### HER2

The established assessment and scoring system for DAKO Herceptest was used^18^. Cases were scored 0, 1+, 2+ or 3+ based on intensity grade of membrane staining and percentage of tumor cells with such staining. 0 and 1 + were classified as negative, and 3 + as positive. HER2 SISH was performed on IHC 2+ cases (Ventana INFORM HER2 DNA probe staining). The 2+ cases were considered HER2 positive if the HER2/Chr17 ratio by SISH was ≥ 2.0^16^ ^19^.

##### Ki67

TMA slides were evaluated for Ki67-scoring, following the approach used on Ki67 evaluation in the Bergen cohort-2 by Knutsvik et al.^16^. For inter-observer assessment, 20 cases in both Bergen cohort-1 and -2 were evaluated by AAS, and thereafter reevaluated by an experienced breast pathologist (GK), who originally assessed Ki67 in the Bergen cohort-2. Both researchers were blinded to patient characteristics. The inter-observer agreement between assessment of Ki67 was excellent (Spearman’s ρ=0.95, kappa-value=1.00). In the inter-observer assessment, GK’s counts were on average 1,3% higher than AAS’ counts. The Ki67 scoring from the Bergen cohort-1 was then corrected using a ratio of 1.013.

##### CK5/6

For CK5/6 a score index (SI) was calculated by multiplying the fraction of tumor cells stained (0: no staining, 1: <10%, 2: 10-50%, 3: >50%) with a score for the staining intensity (0-3). The SI ranged 0-9, where 0 was considered negative, and all other cases positive.

#### Classification of molecular subtypes

The classification of molecular subtypes was done according to St Gallen guidelines 2013^20^, with minor modifications. Luminal A: ER and/or PR positive, HER2 negative, Ki67 low; Luminal B/HER2 negative: ER and/or PR positive, HER2 negative and Ki67 high; Luminal B/HER2 positive: ER and/or PR positive, HER2 positive; HER2 positive (non-luminal): ER and PR negative, HER2 positive; Triple negative: ER, PR, HER2 negative. To dichotomize the Ki67 counts as low/high, the Ki67 cut-off value was derived from the 65th percentile value in hormone receptor (ER and/or PR) positive, HER2 negative cases^21^, calculated as 8.7 % for TMA and 20.4 % for whole sections, including data from both the Bergen cohort-1 and -2.

### Gene expression analyses

#### mRNA cohorts

Two data sets from The Molecular Taxonomy of Breast Cancer International Consortium (METABRIC) were downloaded and analyzed (invasive breast cancer; discovery and validation cohorts; n=997 and n=995 cases, respectively)^22^. The molecular subtypes were defined by the PAM50 classification^23^. Normal breast-like subtypes were excluded from the analyses. Multiple microarray probes covering the same gene were collapsed according to max probe, and log2 transformed data was used in the analyses^22^ ^24^.

#### Gene expression signatures

The luminal progenitor and mature luminal signatures presented by Lim et al.^25^, and two signatures reflecting stemness features^26^ ^27^ were mapped to the METABRIC data. The corresponding signature scores were calculated as described in the primary publications, and if not originally specified, by a sum of expression values of signature genes.

### Statistical analyses

Statistical analyses were performed using IBM SPSS Statistics for Windows, Version 25.0. (Armonk, NY: IBM Corp). A two-sided P value less than 0.05 was considered as statistically significant. Categories were compared using Pearson’s or Fisher’s Exact tests when appropriate. Non-parametric correlations were tested by Spearman’s rank correlation, while Mann-Whitney U and Kruskal-Wallis tests were used to compare continuous variables across groups. Wilcoxon Signed-Rank test was used to compare the differences between two continuous variables. Odds ratios (OR) and their 95% confidence interval were calculated by Mantel-Haenszel method. Kappa statistics were used to test inter- and intra-observer agreement of categorical data. For survival analyses, the endpoint was breast cancer specific survival, defined as the time in months from the date of diagnosis to breast cancer related death. Univariate survival analysis (Kaplan-Meier method) was performed using the log-rank test to compare differences in survival time between categories. Patients who died of other causes or who were alive at the last follow-up, were censored in the analyses. Univariate breast cancer specific survival analysis for Ki67 as continuous variable was performed by Cox’ proportional hazards regression model. When categorizing continuous variables, we applied cut-off points based on median or quartile values, also considering the distribution profile, the size of subgroups, and number of events in survival analyses. See information on separate cut-point for Ki67 in Methods; Molecular subtypes. Only TMA slides (n= 767) was used when Ki67 was analyzed as a continuous variable. For analyses where Ki67 was dichotomized (high, low), information from both TMA and WS (n = 883) was used.

## Results

### Young age relates to aggressive breast cancer features

To elucidate whether there is an age-related distribution of tumor features in young breast cancer patients outside of the mammography screening program, we assessed how clinico-pathologic variables were distributed in patient groups aged below 40 years and 40-49 years at time of diagnosis (Bergen cohort-1). Cases with locally advanced tumors or distant metastases at time of diagnosis were included in the analyses.

The patients presented with age range 25-49 years at time of diagnosis (median 44 years) and the following age distribution: <30: n=10 (3%), 30-39: n=80 (22%), 40-49: n=265 (75%). Age at diagnosis below 40 years was associated with a higher proportion of histologic subtypes other than infiltrating ductal or lobular carcinomas, including 9/90 (10%) cases with medullary carcinomas. Tumor features indicating aggressive breast cancer, including high histologic grade, lymph node metastases, locally advanced disease, ER and PR negative, and HER2 positive status, were more frequent in patients below 40 years (**Supplementary Table 1**).

Next, we compared data from patients in the Bergen cohort-1 (below 50 years at diagnosis) and the Bergen cohort-2 (50-59 years at diagnosis; part of the Norwegian breast cancer screening program, BreastScreen Norway). Patients with distant metastases discovered at time of, or within six months after the primary breast cancer diagnosis, were excluded from these comparative analyses.

Breast cancer patients below 40 years at diagnosis presented more frequently with high histologic grade, lymph node metastases, ER and PR negativity, and HER2 positive status, compared to all other age groups, also when including information on detection mode (screening/interval) in the age group 50-69 years (**Table 1**). To note, we found similar tumor feature frequencies in patients in the age groups 40-49 years and 50-69 years, interval detected (**Table 1**). As expected, patients aged 50-69 years with screening detected tumors, presented with smaller tumors and less frequent aggressive tumor features (**Table 1**).

**Table 1.** Age at diagnosis, detection mode and clinico-pathologic data. Bergen combined cohort (n=890)

|  | Age at diagnosis (years) |  |  |  |  |  |  |  | P |
| --- | --- | --- | --- | --- | --- | --- | --- | --- | --- |
|  | <40 |  | 40-49 |  | 50-69 INT |  | 50-69 SCR |  |  |
|  | n (%) |  | n (%) |  | n (%) |  | n (%) |  |  |
| **Histological type** |  |  |  |  |  |  |  |  |  |
| Ductal | 67 | (77.9) | 225 | (86.9) | 111 | (79.3) | 343 | (85.1) | 0.008 |
| Lobular | 6 | (7.0) | 21 | (8.1) | 21 | (15.0) | 34 | (8.4) |  |
| Other | 13 | (15.1) | 13 | (5.0) | 8 | (5.7) | 26 | (6.5) |  |
| **Histological grade** |  |  |  |  |  |  |  |  |  |
| Low (grade 1 and 2) | 50 | (63.3) | 175 | (71.1) | 102 | (72.9) | 351 | (87.1) | <0.001 |
| High (grade 3) | 29 | (36.7) | 71 | (28.9) | 38 | (27.1) | 52 | (12.9) |  |
| **Tumor diameter** |  |  |  |  |  |  |  |  |  |
| ≤2.0 cm | 48 | (56.5) | 140 | (54.7) | 77 | (55.0) | 337 | (83.6) | <0.001 |
| >2.0 cm | 37 | (43.5) | 116 | (45.3) | 63 | (45.0) | 66 | (16.4) |  |
| **Nodal status** |  |  |  |  |  |  |  |  |  |
| Negative | 36 | (44.4) | 143 | (55.6) | 81 | (59.6) | 314 | (78.3) | <0.001 |
| Positive | 45 | (55.6) | 114 | (44.4) | 55 | (40.4) | 87 | (21.7) |  |
| **Locally advanced** |  |  |  |  |  |  |  |  |  |
| No | 71 | (82.6) | 237 | (90.8) | 121 | (86.4) | 387 | (78.3) | <0.001 |
| Yes | 15 | (17.4) | 24 | (9.2) | 19 | (13.6) | 15 | (3.7) |  |
| **ER** |  |  |  |  |  |  |  |  |  |
| Positive | 37 | (43.5) | 188 | (72.0) | 99 | (70.7) | 361 | (89.6) | <0.001 |
| Negative | 48 | (56.5) | 73 | (28.0) | 41 | (29.3) | 42 | (10.4) |  |
| **PR** |  |  |  |  |  |  |  |  |  |
| Positive | 36 | (42.4) | 192 | (73.6) | 85 | (60.7) | 301 | (74.7) | <0.001 |
| Negative | 49 | (57.6) | 69 | (26.4) | 55 | (39.3) | 102 | (25.3) |  |
| **HER2** |  |  |  |  |  |  |  |  |  |
| Negative | 51 | (65.4) | 220 | (88.7) | 117 | (83.6) | 345 | (88.0) | <0.001 |
| Positive | 27 | (34.6) | 28 | (11.3) | 23 | (16.3) | 47 | (12.0) |  |
| **Ki67 (combined)** |  |  |  |  |  |  |  |  |  |
| Low | 25 | (32.9) | 127 | (51.2) | 65 | (46.4) | 256 | (65.5) | <0.001 |
| High | 51 | (67.1) | 121 | (48.8) | 75 | (53.6) | 135 | (34.5) |  |
| **Molecular subtypes (IHC)** |  |  |  |  |  |  |  |  |  |
| Luminal A | 16 | (20.5) | 109 | (44.3) | 61 | (43.6) | 233 | (59.6) | <0.001 |
| Luminal B/HER2 neg | 15 | (19.2) | 75 | (30.5) | 34 | (24.3) | 85 | (21.7) |  |
| Luminal B/HER2 pos | 10 | (12.8) | 16 | (6.5) | 9 | (6.4) | 37 | (9.5) |  |
| HER2 positive | 17 | (21.8) | 12 | (4.9) | 14 | (10.0) | 10 | (2.6) |  |
| Triple negative | 20 | (25.6) | 34 | (13.8) | 22 | (15.7) | 26 | (6.6) |  |
| **CK5/6** |  |  |  |  |  |  |  |  |  |
| Negative | 63 | (76.8) | 217 | (85.4) | 114 | (81.4) | 352 | (89.6) | 0.007 |
| Positive | 19 | (23.2) | 37 | (14.6) | 26 | (18.6) | 41 | (10.4) |  |
| **Basal-like phenotype\*** |  |  |  |  |  |  |  |  |  |
| Negative | 74 | (88.1) | 234 | (91.8) | 126 | (90.0) | 386 | (95.8) | 0.016 |
| Positive | 10 | (11.9) | 21 | (8.2) | 14 | (10.0) | 17 | (4.2) |  |

n: number of patients; INT: interval detected; SCR: screen detected; P-values by Pearson's chi-square test. \* Basal-like phenotype defined as triple negative and CK5/6 positive. Missing data: Histological type: n=22; tumor diameter: n=6; nodal status: n=15; ER: n=1; PR: n=1; HER2: n=32; Ki67: n=41 (Ki67 combined is data from TMA and from WS when TMA was missing); Molecular subtypes: n=35; CK5/6: n=28; Basal-like phenotype: n=38.

The molecular subtypes were determined by the IHC markers ER, PR, HER2 and Ki67 (for details, see Methods). Age below 40 years was associated with more frequent HER2 and triple negative subtypes (**Table 1; Figure 1A**).

**Figure 1.**
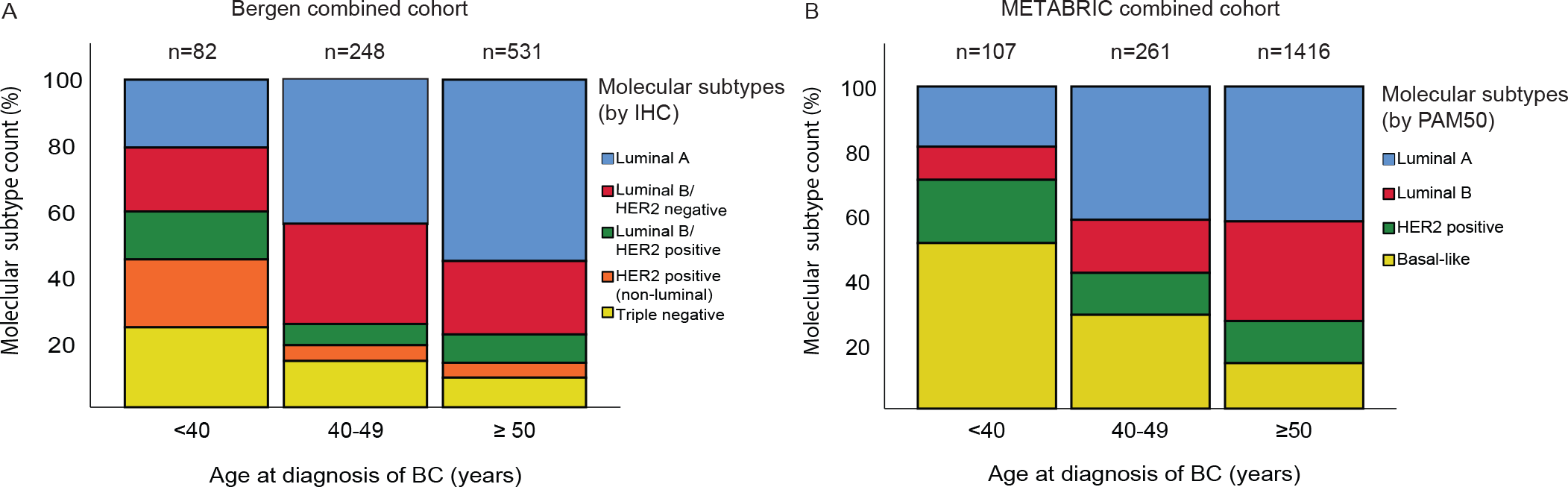
**A-B**. Distribution of molecular subtypes by % of cases within each age groups. A) Bergen combined breast cancer cohort (n=861). Molecular subtypes defined by IHC markers. B) METABRIC combined cohort (Discovery + Validation, n=1784). Molecular subtypes defined by PAM50.

**Figure 1.**
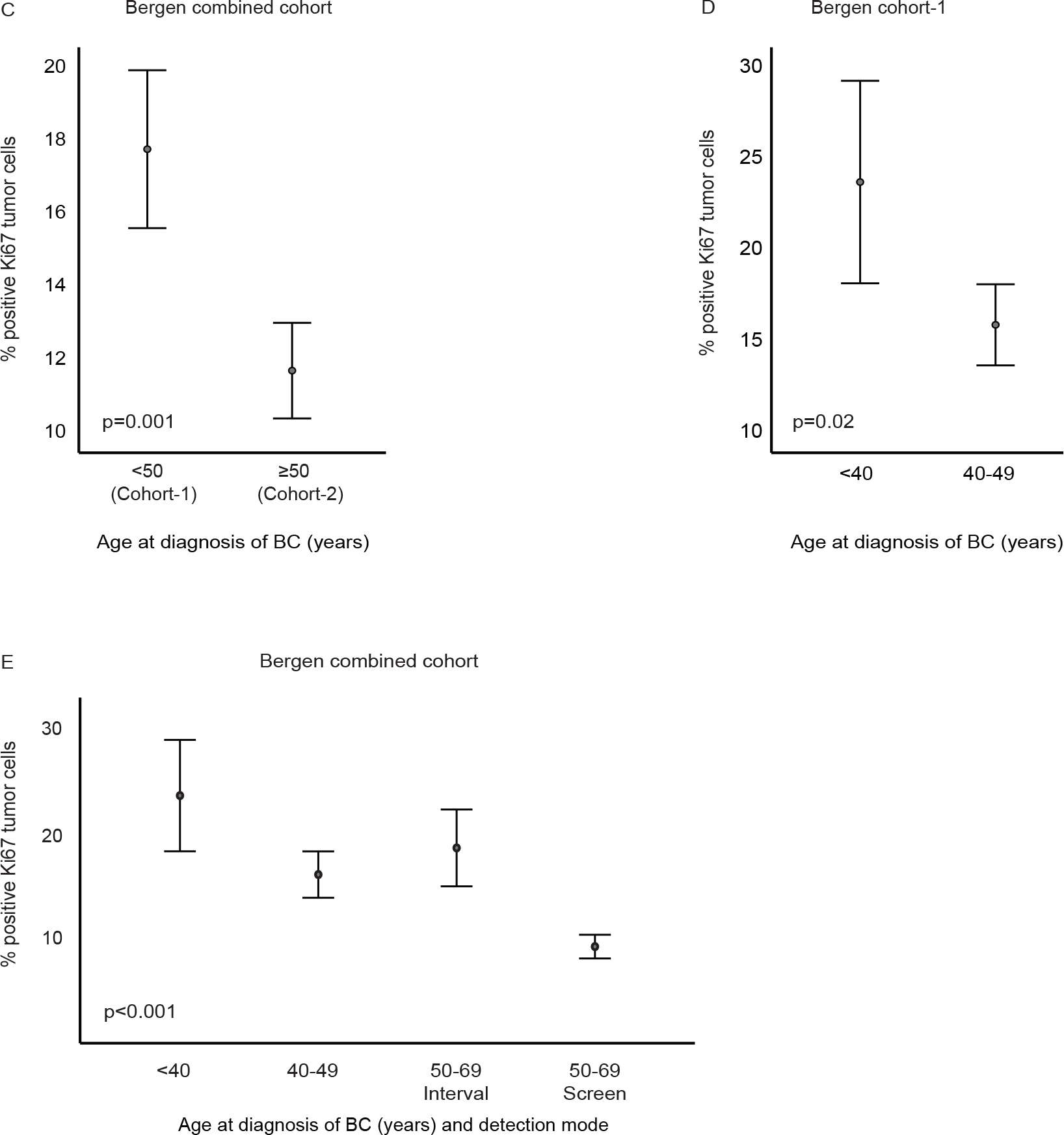
**C-E.** Ki67 positive tumor cells (%) from TMA across age groups and detection mode. Data is presented by error-bars with 95% confidence interval of the mean, and p-values by the Mann-Whitney U test (C-D) or Kruskal-Wallis test (E). C, E) Bergen combined cohort (n=774). D) Bergen cohort-1 (n=318).

In data from the combined METABRIC cohorts (n= 1784, after excluding cases with the normal breast-like subtype), we validated that breast cancer at age below 40 years was associated with higher histologic grade, more frequent lymph node metastases, and ER negative tumors (**Supplementary Table 2**). The ESR1 (encoding ERα) mRNA expression was lower in breast cancer tissue from patients below 40 years (**Supplementary Figure 2**), in line with the observations of more frequent ER negative tumors in this patient subset.

### Higher tumor cell proliferation in breast cancer of the young

For the combined Bergen cohorts (n=767), Ki67 ranged from 0.0-86.5% tumor cell positivity (TMA-based assessment). Mean Ki67 was 14.3%, and median 8.1% (Bergen cohort-1 and -2, combined). Tumors from patients in the Bergen cohort-1 demonstrated higher Ki67 levels compared to the Bergen cohort-2; median Ki67: 10.5% and 7.0%, respectively (P≤0.001; **Figure 1C**). We found higher Ki67 levels in patients below 40 years compared to the other age groups (**Figure 1D-E; Table 1**).

When stratifying the analyses by molecular subgroups, the luminal B/HER2 negative tumors showed higher Ki67 scoring in the young (median 16.8% in the Bergen cohort-1 and 13.6% in the Bergen cohort-2; **Supplementary figure 3A**). Luminal B/HER2 negative tumors also showed higher Ki67 in patients aged 40-49 years when compared to patients aged 60-69 years (P=0.025), and when compared to patients aged 50-59 years (P=0.045; **Supplementary figure 3B**). In the Bergen cohort-1, high levels of Ki67 were associated with larger tumor diameter, high histologic grade, ER and PR negativity, HER2 positivity, and the molecular subtypes except luminal A (**Table 2**).

**Table 2.** Ki67, clinico-pathologic variables and molecular subtypes. Bergen breast cancer cohort-1 (age <50 years at time of diagnosis, n=347).

|  | % Ki67 positive tumor cells* |  | OR (95% CI) | P |
| --- | --- | --- | --- | --- |
|  | Low<br>n (%) | High<br>n (%) |  |  |
| **Age at diagnosis** |  |  |  |  |
| 40-49 years | 109 (46.0) | 128 (54.0) | 1.0 | 0.023 |
| < 40 years | 23 (31.1) | 51 (68.9) | 1.9 (1.1-3.3) |  |
| **Histological grade** |  |  |  |  |
| Low (grade 1 and 2) | 112 (54.9) | 92 (45.1) | 1.0 | <0.001 |
| High (grade 3) | 12 (13.2) | 79 (86.8) | 8.6 (5.5-13.5) |  |
| **Tumor diameter** |  |  |  |  |
| ≤2.0 cm | 81 (47.4) | 90 (52.6) | 1.0 | 0.06 |
| >2.0 cm | 50 (36.8) | 86 (63.2) | 2.05 (1.5-2.8) |  |
| **Nodal status** |  |  |  |  |
| Negative | 77 (47.8) | 84 (52.2) | 1.0 | 0.07 |
| Positive | 54 (37.5) | 90 (62.5) | 1.5 (0.97-2.4) |  |
| **ER** |  |  |  |  |
| Positive | 100 (48.5) | 106 (51.5) | 1.0 | 0.002 |
| Negative | 32 (30.5) | 73 (69.5) | 3.6 (2.5-5.2) |  |
| **PR** |  |  |  |  |
| Positive | 105 (50.5) | 103 (49.5) | 1.0 | <0.001 |
| Negative | 27 (26.2) | 76 (73.8) | 2.9 (1.7-4.8) |  |
| **HER2** |  |  |  |  |
| Negative | 118 (45.7) | 140 (54.3) | 1.0 | 0.003 |
| Positive | 12 (23.5) | 39 (76.5) | 4.2 (2.6-6.6) |  |
| **Molecular subtypes (IHC)** |  |  |  |  |
| Luminal A | 107 (100) | 0 (0) |  | <0.001 |
| Luminal B/HER2 neg | 0 (0) | 99 (100) |  |  |
| Luminal B/HER2 pos | 6 (25) | 18 (75.0) |  |  |
| HER2 positive | 6 (22.2) | 21 (77.8) |  |  |
| Triple negative | 11 (21.2) | 41 (78.8) |  |  |
| **CK5/6** |  |  |  |  |
| Negative | 125 (48.6) | 132 (51.4) | 1.0 | <0.001 |
| Positive | 7 (13.0) | 47 (87.0) | 6.4 (2.8-14.6) |  |
| **Basal-like phenotype\*\*** |  |  |  |  |
| Absent | 130 (46.9) | 147 (53.1) | 1.0 | <0.001 |
| Present | 2 (6.1) | 31 (93.9) | 13.7 (3.2-58.4) |  |

n: number of patients; P-values by Pearson's chi-square test. \* Ki67 count per TMA; cut off 8.7% (65th percentile in HER2 negative luminal tumors). \*\* Basal-like phenotype defined as triple negative and CK5/6 positive. Missing data: Tumor-diameter: n=40; Histological grade: n=52; Lymph node status: n=42; ER: n=36; PR: n=36; HER2: n=38; Ki67: n=36; Molecular subtypes: n=38; CK5/6: n=36; Basal-like phenotype: n=39

### Young age is associated with a basal-like phenotype and features of stemness

Due to the strong association seen between young age and the triple negative phenotype, we next asked how age is related to the basal-like phenotype. In the Bergen cohorts, age below 40 years was associated with basal-like differentiation, both as defined by CK5/6 tumor cell positivity alone, and concurrent triple negative subtype and CK5/6 positivity (**Table 1**). Consistent with this, young age was significantly associated with the basal-like subgroup (by PAM50) in the METABRIC cohort (**Supplementary Table 2; Figure 1B**).

Following up on the association between young age and a basal-like phenotype, we assessed how transcriptional patterns of mammary stem cells and luminal progenitor cells present in different age groups. Age below 40 years was significantly associated with a luminal progenitor signature score^25^ and two additional signature scores reflecting cancer stem cell activation^26^ ^27^, **(Figure 2A-C).** A mature luminal signature score was lower expressed in breast cancer of the young (**Figure 2D**).

**Figure 2.**
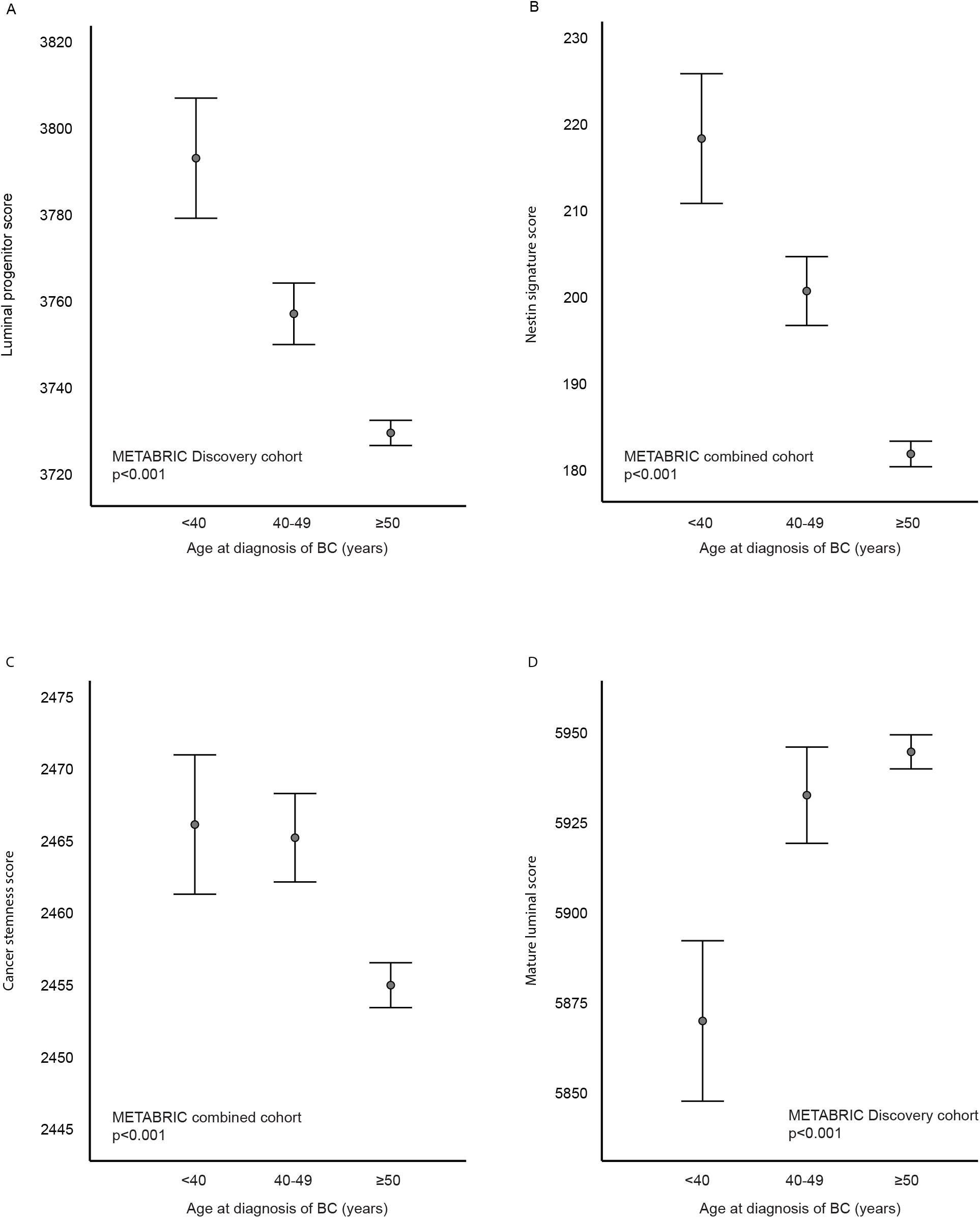
Gene expression signatures reflecting progenitor and stemness features across age groups in METABRIC cohort (Discovery n=997, combined n=1992). Higher scores of signatures reflecting A) Luminal progenitor activation; B-C) stemness features in breast cancer of the young. D) Lower scores reflecting a mature luminal program in the young. Data is presented by error-bars with 95% confidence interval of the mean, and p-values by Kruskal-Wallis test.

The association between higher cancer stem cell transcriptional patterns and young age was also present within each of the PAM50-defined molecular subtypes. Patients below 40 years showed a higher stemness-related Nestin score in luminal A and basal-like subtypes when compared to patients older than 50 years (P≤0.001; **Supplementary Figure 4A**). For all subtypes, the Nestin score was higher in patients aged 40-49 years when compared to patients older than 50 years (P≤.042; **Supplementary Figure 4A**). For the cancer stemness score, the difference was restricted to the luminal subtypes. In luminal A, both patient groups <40 and 40-49 presented higher cancer stemness score when compared to patients older than 50 years (P=0.011 and P<0.001, respectively). In luminal B tumors in patients aged 40-49 years, the cancer stemness score was significantly higher than those older than 50 years (P=0.001; **Supplementary Figure 4B**). The mature luminal score was lower in patients aged 40-49 with luminal B tumors when compared to patients older than 50 (P=0.035; **Supplementary Figure 4C**), while there was no difference between subtypes for the luminal progenitor score (**Supplementary Figure 4D**). Taken together, these results support a link between breast cancer of the young, a basal-like phenotype, and tumors with progenitor and stem cell features.

### Shorter survival in breast cancer of the young

Based on our results, indicating a more aggressive tumor biology in the young breast cancer patients, we next assessed the age-related disease-specific survival patterns. Patients in the age groups below 40 years; 40-49 years; 50-59 years; 60-69 years, demonstrated declining survival, from the older age groups with the longest disease-specific survival, to patients below 40 years showing poorest survival (Bergen combined cohorts; **Figure 3A**).

**Figure 3.**
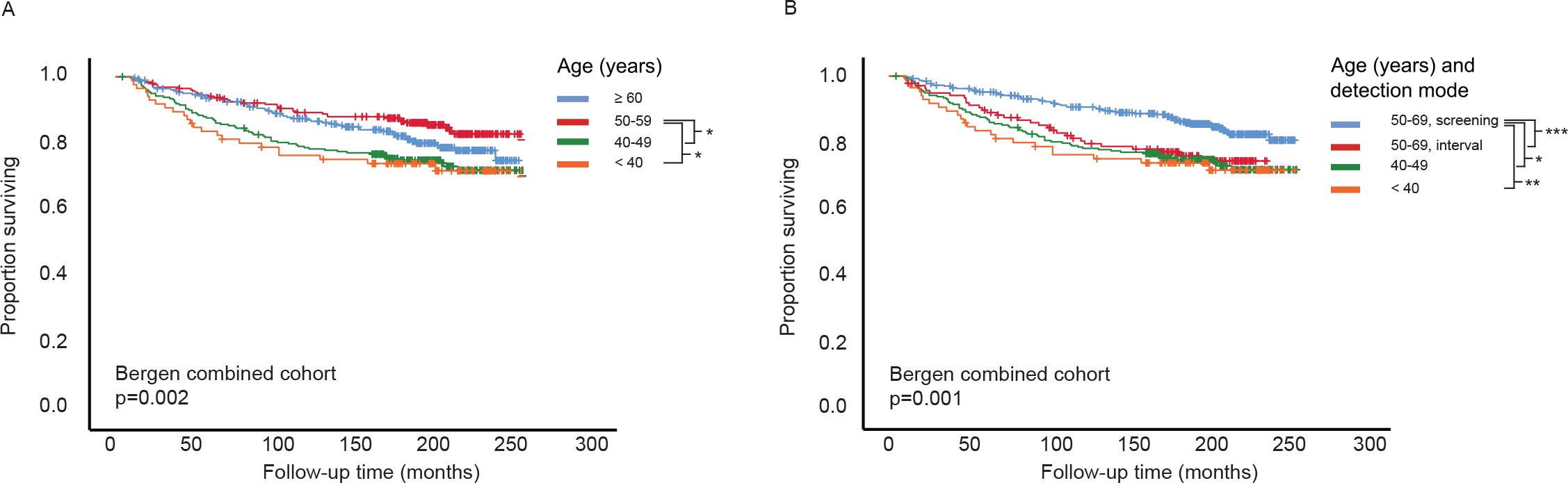
**A-B**. Breast cancer disease specific survival according to age at diagnosis and detection mode for age 50-69 (screening or interval detected). Kaplan-Meier univariate breast cancer specific survival analysis (log-rank test for difference) in Bergen breast cancer cohort (cohort-1 and -2, n=898). A) Age groups <40 years; 40-49 years; 50-59 years; ≥60 years at time of diagnosis. B) Age groups <40 years; 40-49 years; 50-69 years, including information on detection mode for the age group 50-69.

**Figure 3.**
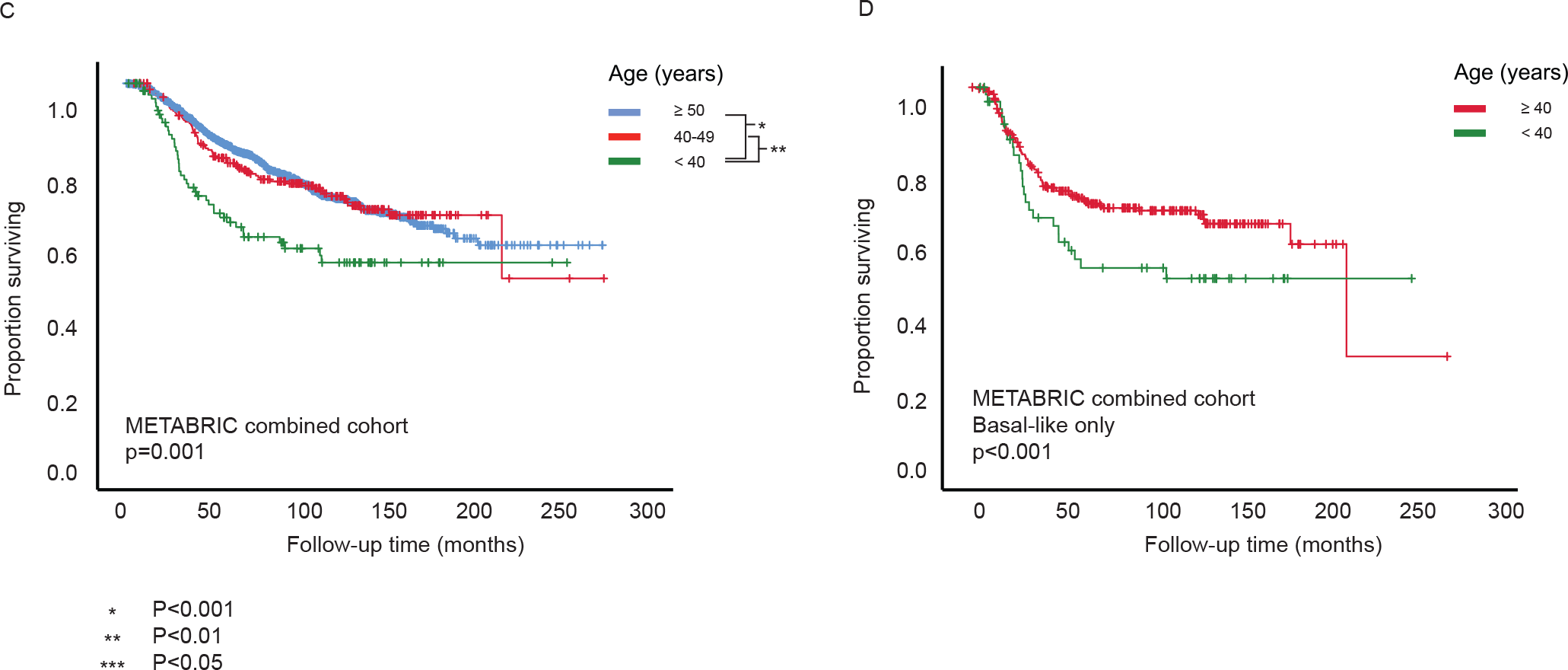
**C-D.** Kaplan-Meier univariate breast cancer disease specific survival analysis in the METABRIC combined (Discovery + Validation, n=1990) cohort according to age groups (log-rank test for difference). C) Age groups <40 years; 40-49 years; ≥50 years, at time of diagnosis. D) Basal-like only; age groups <40 years; ≥40 years, at time of diagnosis.

When adding information about mode of tumor detection, *i.e.* screening or interval detection, and detection outside of the breast cancer screening program, the 5-year disease-specific survival was declining from 95% in the older age groups (50-69 years) diagnosed by screening, through 89.1 % for the older age group diagnosed between screening rounds (interval cancer), and 87.3% for the age group 40-49 years, to poorest 5-year survival of 82.5% at age below 40 at time of diagnosis (Bergen combined cohorts; **Figure 3B**).

In the METABRIC cohort, patients below age 40 at time of diagnosis validated significantly shorter disease-specific survival compared to patients aged 40-49 years and ≥ 50 years (**Figure 3C**). When stratifying by molecular subtypes, age-dependent survival analyses demonstrated shorter disease-specific survival in patients below 40 years only in the basal-like subgroup (**Figure 3D**).

Taken together, our data demonstrate age-related clinical impact with respect to survival, paralleling the age-related phenotypic alterations seen in breast cancer in young patients.

### Age dependent prognostic value of Ki67

In the Bergen cohort-1, high Ki67 tumors were associated with shorter disease-specific survival (**Figure 4A**). When combining Ki67 information from both in-house cohorts in survival analyses (Bergen combined cohort-1 and -2; n=883; combined WS and TMA, see Methods), Ki67-high tumors were associated with shorter disease-specific survival in patients below 50 years (Bergen cohort-1) compared to patients ≥50 years (Bergen cohort-2) (P=0.044; **Figure 4B**). This indicates added value when including age to the prognostication in breast cancer. Next, aiming to see whether different age-groups presented with distinct Ki67 cut-points providing the strongest prognostic value, we analyzed Ki67 categorized into high vs low using percentile-based cut-points. However, none of the cut-points showed statistically significant prognostic value in the age group below 40. On the other hand, for all other age groups (including screening/interval separation), a statistically significant prognostic value was identified for one or several different Ki67 cut-points (**Supplementary Table 3**), indicating a weaker value for Ki67 as prognosticator in breast cancer patients below 40 years. The results were confirmed when redoing the univariate survival analyses with the Ki-67 scoring as a continuous variable (Cox’ proportional hazards regression model; **Supplementary Table 3**).

**Figure 4.**
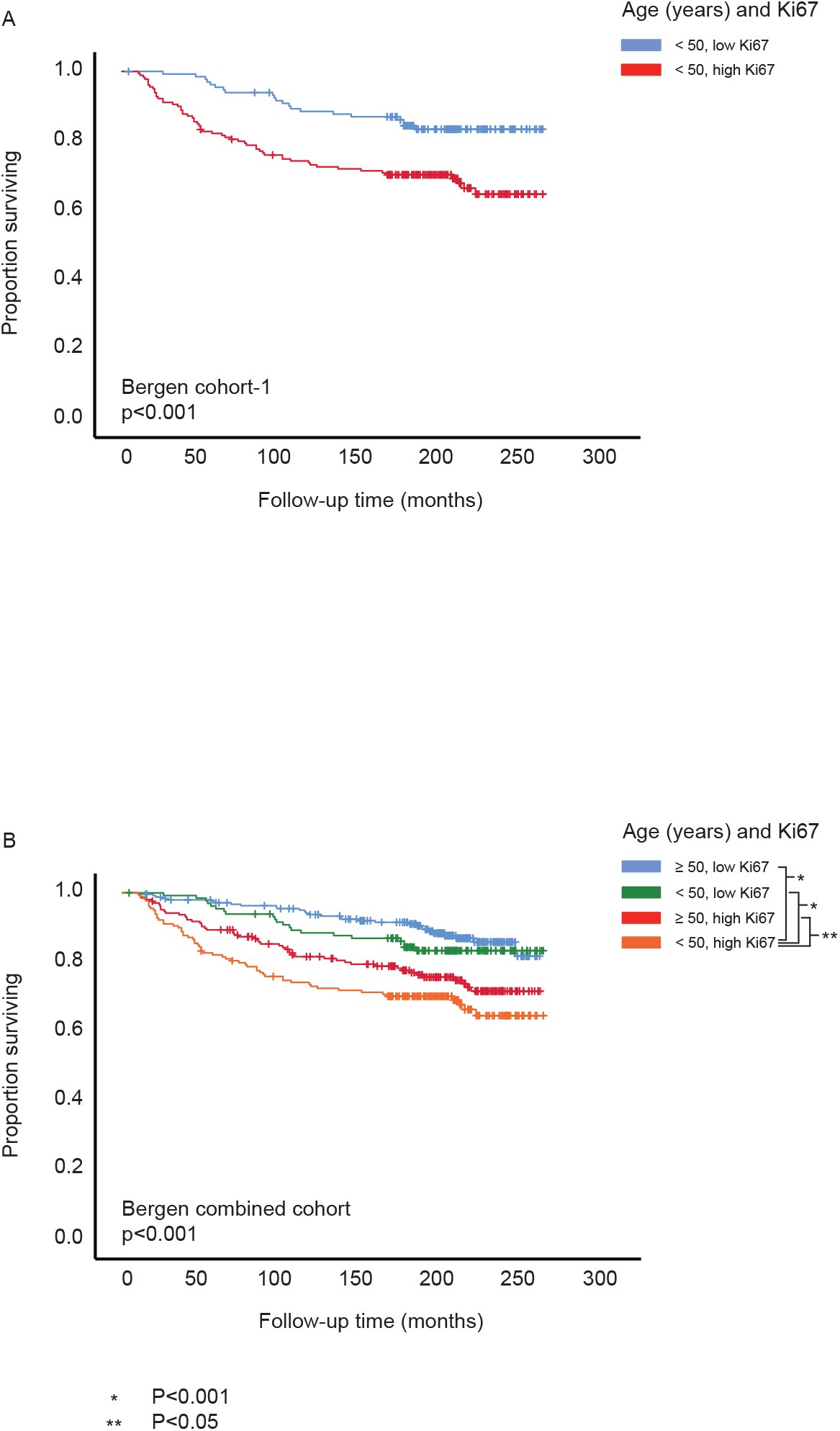
Breast cancer disease specific survival according to age at diagnosis and Ki67. Kaplan-Meier univariate breast cancer specific survival analysis (log-rank test for difference) in the Bergen Breast cancer cohort. A) Cohort-1 (<50 years at diagnosis, n=320); B) Bergen Cohort-1 and -2 (n=843).

Summarized, our data supports an age-related distribution of breast tumor features impacting clinical outcome, and aggressive tumor characteristics more frequently presented in patients below 40 years. We show increased proliferation by Ki67 in tumors of the young. Survival analyses indicate added value by including age to the prognostication, and a weaker prognostic value of Ki67 in the young.

## Discussion

Previous studies have shown that breast cancer in young women have a poorer prognosis and a higher proportion of aggressive subtypes compared to older age groups^6^ ^7^ ^9^ ^28^. However, there is an ongoing debate whether the worse prognosis seen in younger women is due to unique biology or age-specific variations in subtypes alone. In line with previous studies on breast cancer of the young^29^, we confirm an overrepresentation of aggressive tumor characteristics among the younger patients. To the best of our knowledge, this is the first population-based study to investigate phenotypic profiles and survival in breast cancer of the young compared to screening or interval detected cancers in older patients.

We demonstrated the strongest association with aggressive tumor features in the patient group below 40. The age group 40-49 and the interval detected cancers in patients aged 50-69 showed a similar distribution of aggressive tumor characteristics, and the screening detected cancers in patients aged 50-69 presented the least aggressive tumor features. Interval detected cancers have previously been shown to have a more aggressive phenotype than screening detected cancers^30^. Due to a lack of mammography screening for the women below 50 years, it is not surprising that the younger breast cancer patients show a more aggressive course than the screening-detected cancers. However, the identified separation of the clinico-pathologic characteristics between patients below 40 and those aged 40-49, points to age-related phenotypes beyond the differences due to screening/interval detection. To note, we do not have information regarding *BRCA1* or *2* statuses for either Bergen cohorts. Higher prevalence of mutated *BRCA* genes in younger women will contribute to the increased aggressive tumor characteristics seen in this patient-group^31^.

In our study, the younger patients (below 40) present overall higher Ki67 scoring compared to older patients, also within the Luminal B subtype. Corresponding to this, previous studies have shown age-related differences for Ki67 also within molecular subtypes – with patients below 40 years presenting higher Ki67 in Luminal A, Luminal B, Luminal B/HER2 negative, and the triple negative subtypes^7^ ^32^. A recent study from our group analyzed age-related gene expression differences and found overexpression of genes related to proliferation in the young, also in the luminal subsets^33^. Together, this points to tumors of high proliferation in young patients, especially within the luminal subgroups.

A part of our in-house cohort (Bergen cohort-2) was included in the national breast screening program (age 50-69 years). Thus, whereas breast cancer in young women follows a natural clinical course, the screening program may detect asymptomatic tumors with less aggressive tumor features, including low proliferation. This might cause a bias when comparing tumor features in patients above and below 50 years. To account for this, we redid the Ki67 analyses, splitting the Bergen cohort-2 by detection mode, screening vs interval. Younger patients still showed a trend of higher Ki67, across detection mode. However, it was only statistically significant higher Ki67 in the young patients when compared with the older patients of the screening group.

Our results support a link between breast cancer of the young, a basal-like phenotype, and tumors with increased progenitor and stem cell features. In previous studies, mammary stem cells have been proposed with important roles in breast cancer development and progression^34^, and with potential relevance as targets in new treatment strategies^34^ ^35^. Aging is by various mechanisms proposed to stimulate cancer stem cell development^36^. In contrast, our data indicates more frequent cancer stemness in the younger age groups – proposing additional perspectives to the view on aging and cancer stem cells.

Previous studies by Azim *et al*. concluded that young age adds extra biological complexity independent of differences in subtype distribution. They found enrichment of processes related to immature mammary epithelial cells and growth factor signaling, and gene signatures linked to increased proliferation, stem cell features and endocrine resistance, were highly expressed among the young^10^ ^13^. Similarly, Johnson *et al*. showed age-related differences with higher expression of several genes related to tumor proliferation, invasion, and metastasis in the young patients^37^. Our results are in line with these findings, supporting our conclusion to consider the age-related biology in breast cancer care.

In our in-house, population-based cohorts, we demonstrated shorter breast cancer specific survival among the youngest patients (below 40). The same applied for the METABRIC cohort, where age below 40 associated with shorter survival than older age in patients with basal-like tumors. However, a survival effect of age was not found in the other molecular subtypes. Previous studies on breast cancer of the young have demonstrated relations between age and survival in different subsets of tumor subtypes, maybe depending on the composition of the cohorts studied^7^ ^13^ ^38-41^.

Ki67 showed no prognostic significance for overall breast cancer specific survival in patients younger than 40 years. For all other age-groups, several Ki67 cut-points presented with prognostic value. Previous studies have shown similar results, where Ki67 has shown prognostic value for distant disease-free survival in subsets of the young, like PR positive Luminal B/HER2 negative tumors^42^. When applying disease-specific survival as endpoint in our study, Ki67 did not show statistically significant prognostic value in patients below 40, neither when analyzing jointly all subtypes, nor when separated into molecular subgroups. When analyzing only the age group younger than 40, one should keep in mind the potential bias that the smaller group size may affect the statistical power, especially when further categorized in different molecular sub-groups.

In conclusion, our population-based study confirms previous findings of more aggressive breast cancer phenotypes in patients below 40 years at time of diagnosis. We point to increased tumor cell proliferation and stem-like features in breast cancer of the young, with likely clinical relevance. These results support continued search for age-specific treatment and follow-up strategies, including a focus on proliferation and cancer stemness.

## Author contribution

EW conceived and designed the study, and contributed to pathology review of tumor sections, data analysis and interpretation, and writing the manuscript. LAA contributed to the study design, data analyses and interpretation, and to writing the manuscript. AAS contributed to the study design, performed the tissue-based work and immunohistochemistry, performed statistical analysis and data interpretation, and contributed to writing the manuscript. ROCH contributed to data collection, data analyses, results interpretation, and to writing the manuscript. GK participated in data collection and interpretation, and training in Ki67 assessment. AKMS, CA, LMI, UH, ABK, AFT, KK, BD, EAH, TA, and IMS participated in data collection and interpretation. All authors commented on and approved the final manuscript.

## Acknowledgements and funding information

We thank Ingeborg Winge, Gerd Lillian Hallseth, Randi Hope Lavik, and Bendik Nordanger for their excellent technical assistance. This work was partly supported by the Research Council of Norway through its Centre of Excellence funding scheme, project number 223250. The study was also supported by the University of Bergen (Medical Student Research Program), Norwegian Cancer Society and Helse Vest RHF.

## Data availability

Data from the Molecular Taxonomy of Breast Cancer International Consortium (METABRIC) is available at https://ega-archive.org/studies/EGAS00000000083.

