## supplementary files for "Age-related phenotypes in breast cancer: a population-based study"

The supplementary information contains:

Supplementary figure 1: Overview of cases in Bergen breast cancer cohort-1.

Supplementary figure 2: ESR1 (ER) and PGR (PR) mRNA expression across age groups, METABRIC combined cohort.

Supplementary figure 3: Ki67 positive tumor cells (%) across age groups and molecular subtypes.

Supplementary figure 4: Gene expression signatures reflecting progenitor and stemness features across age groups, METABRIC combined cohort.

Supplementary table 1: Age at diagnosis and clinico-pathologic data, Bergen breast cancer cohort-1.

Supplementary table 2: Age at diagnosis, clinico-pathologic data, METABRIC combined cohort.

Supplementary table 3: Age at diagnosis and prognostic value of Ki67 TMA scoring, Bergen combined cohort.

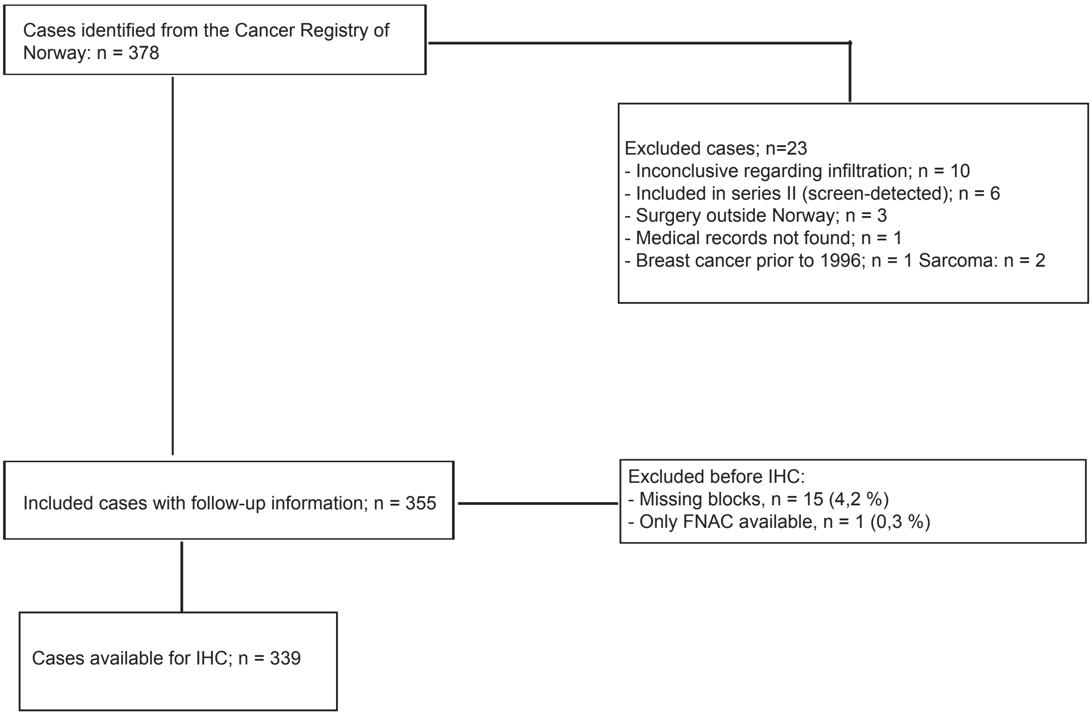


**Supplementary figure 1.** Overview of cases in Bergen breast cancer cohort-1; breast cancer cases diagnosed at age below 50 years at time of diagnosis. n=number of cases; IHC=immunohistochemistry; FNAC=fine needle cytology aspiration.

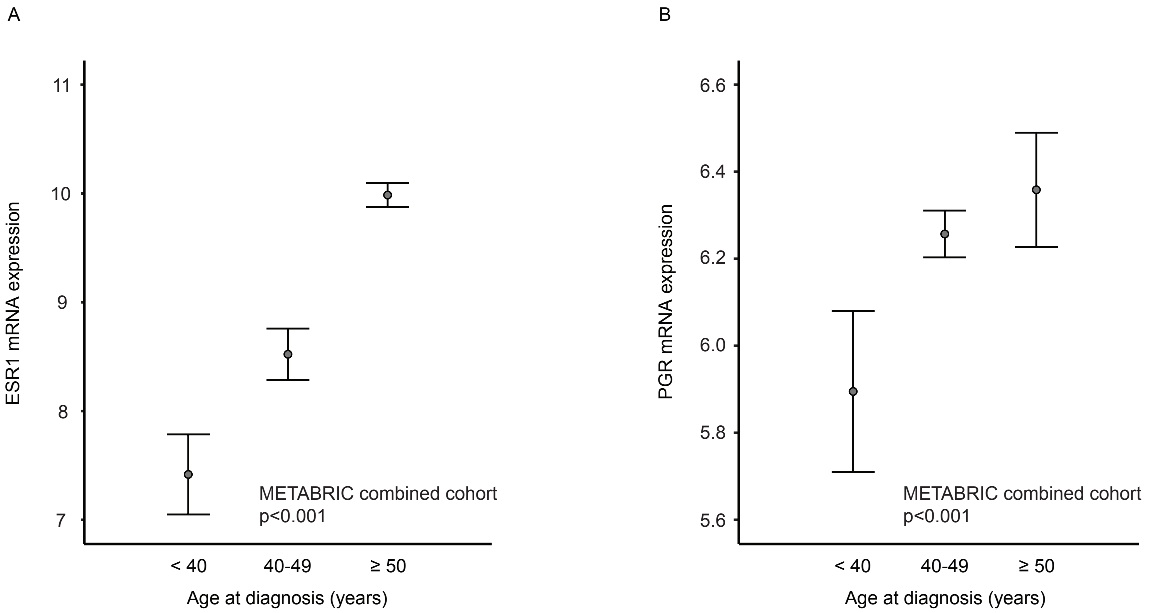


**Supplementary figure 2.** ESR1 (ER) and PGR (PR) mRNA expression across age groups. METABRIC combined cohort (Discovery + Validation, n=1784). Data is presented by error-bars with 95% confidence interval of the mean, and p-values by the Kruskal-Wallis test.

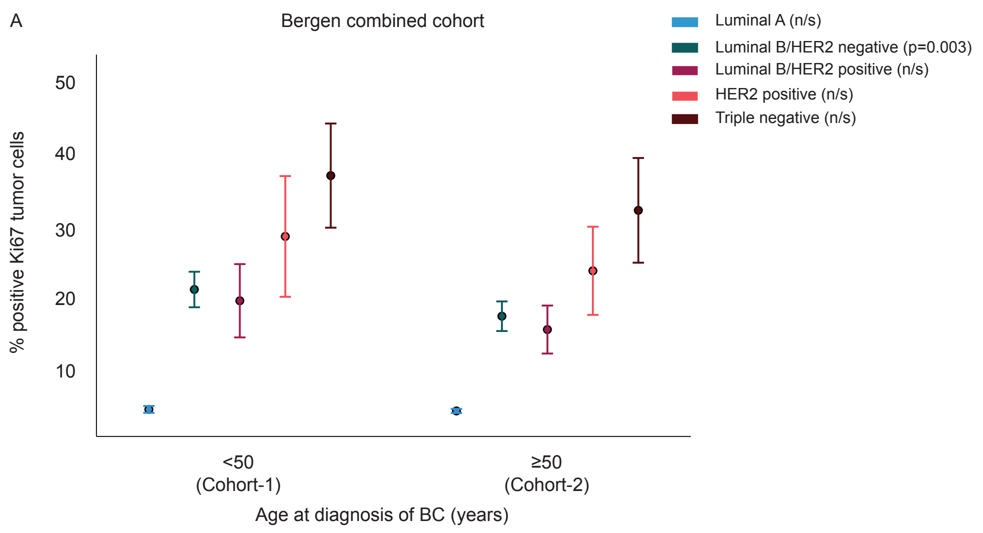


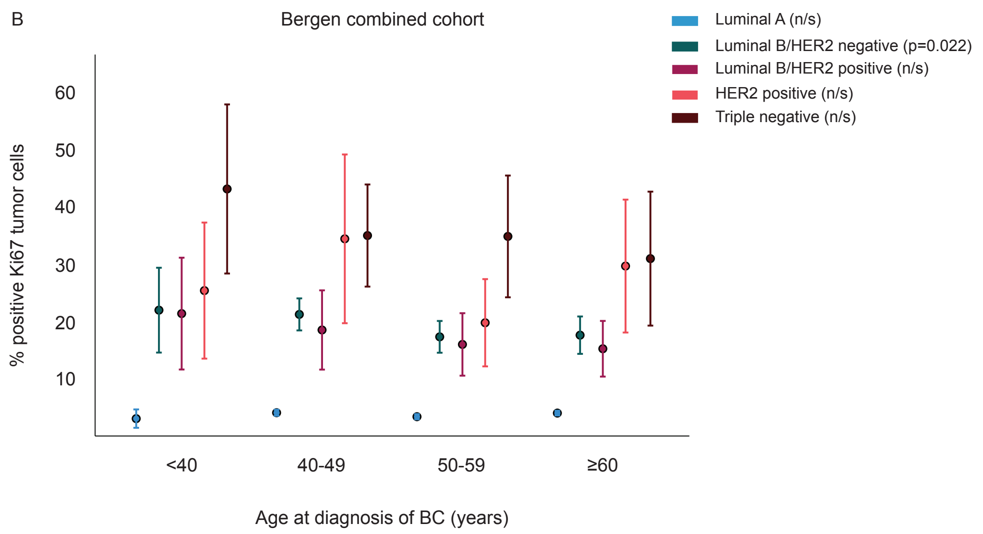


**Supplementary figure 3.** Ki67 positive tumor cells (%) across age groups and molecular subtypes. Bergen combined cohort (n=861). Data is presented by error-bars with 95% confidence interval of the mean, and p-values by the Mann-Whitney U test (A) or Kruskal-Wallis test (B).

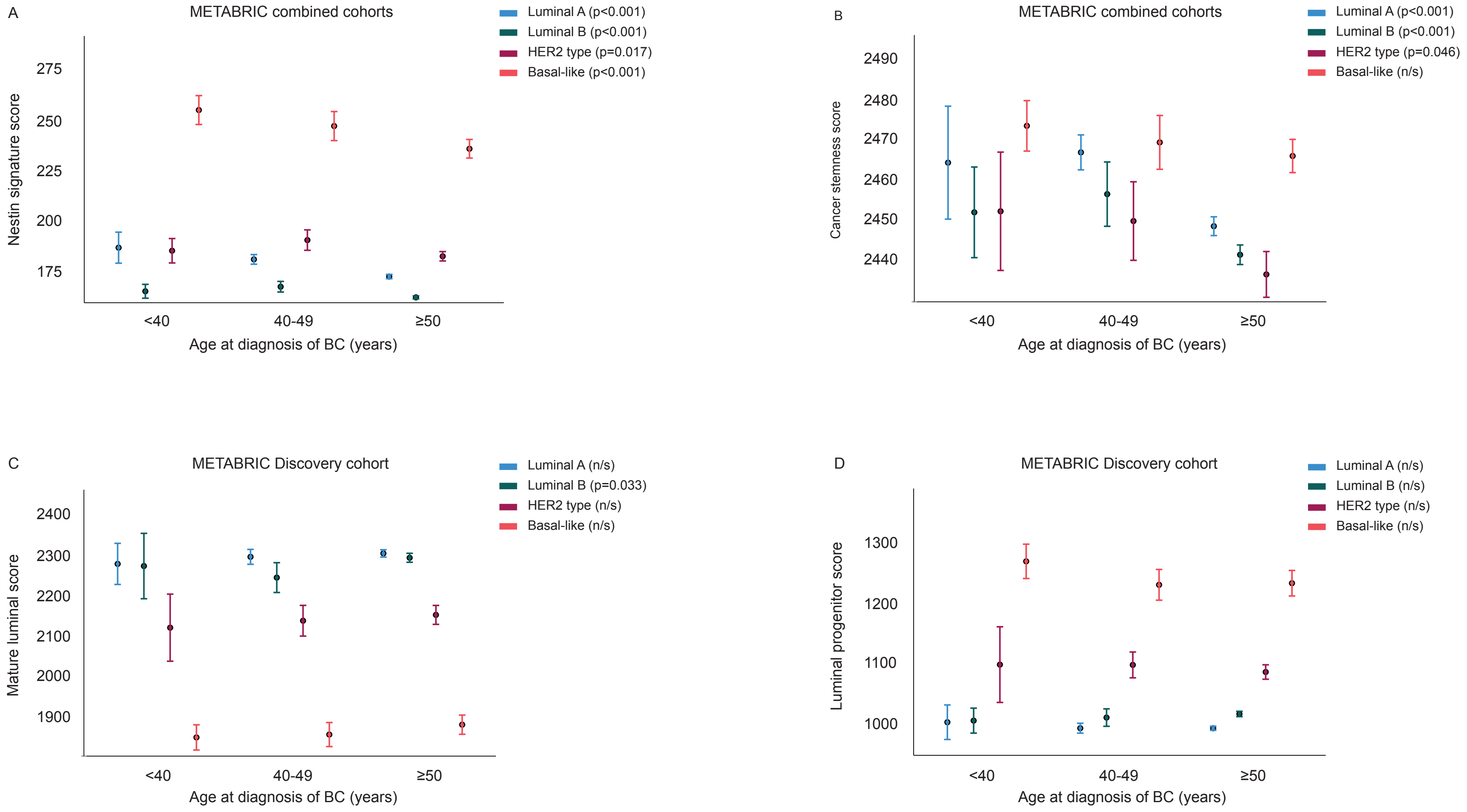


**Supplementary figure 4.** Gene expression signatures reflecting progenitor and stemness features across age groups (METABRIC cohort, Discovery n=997; combined n=1992). Higher scores of the signatures reflecting A-B) stemness features in breast cancer of the young; D) Luminal progenitor activation. C) Lower scores of the signature reflecting a mature luminal program in the young. Data is presented by error-bars with 95% confidence interval of the mean, and p-values by the Kruskal-Wallis test.

**Supplementary table 1.** Age at diagnosis and clinico-pathologic data. Bergen breast cancer cohort-1 (n=355)

| | Age at diagnosis (years) | | | | | |
|---|---|---|---|---|---|---|
| | <40 | | 40-49 | | | |
| | n (%) | | n (%) | | OR (95% CI) | P |
| **Histological type** | | | | | | |
| Ductal | 71 | (78.9) | 228 | (86.7) | | |
| Lobular | 6 | (6.7) | 22 | (8.4) | | 0.011 |
| Other | 13 | (14.4) | 13 | (4.9) | | |
| **Histological grade** | | | | | | |
| Low (grade 1 and 2) | 52 | (63.4) | 177 | (71.1) | 1.0 | 0.215 |
| High (grade 3) | 30 | (36.6) | 72 | (28.9) | 1.4 (0.8-2.4) | |
| **Tumor diameter** | | | | | | |
| ≤2.0 cm | 49 | (55.1) | 141 | (54.2) | 1.0 | 0.903 |
| >2.0 cm | 40 | (44.9) | 119 | (45.8) | 0.97 (0.6-1.6) | |
| **Nodal status** | | | | | | |
| Negative | 36 | (42.4) | 143 | (55.0) | 1.0 | 0.046 |
| Positive | 49 | (57.6) | 117 | (45.0) | 1.7 (1.01-2.7) | |
| **Locally advanced** | | | | | | |
| No | 73 | (81.1) | 240 | (90.6) | 1.0 | 0.023 |
| Yes | 17 | (18.9) | 25 | (9.4) | 2.2 (1.1-4.4) | |
| **Distant metastasis** | | | | | | |
| No | 86 | (95.6) | 261 | (98.5) | 1.0 | 0.11 |
| Yes | 4 | (4.4) | 4 | (1.5) | 3.0 (0.7-12.4) | |
| **ER** | | | | | | |
| Positive | 39 | (43.8) | 190 | (71.7) | 1.0 | <0.001 |
| Negative | 50 | (56.2) | 75 | (28.3) | 3.2 (1.98-5.3) | |
| **PR** | | | | | | |
| Positive | 38 | (42.7) | 193 | (72.8) | 1.0 | <0.001 |
| Negative | 51 | (57.3) | 72 | (27.2) | 3.5 (2.1-5.8) | |
| **HER2** | | | | | | |
| Negative | 53 | (64.6) | 222 | (88.8) | 1.0 | <0.001 |
| Positive | 29 | (35.4) | 28 | (12.8) | 4.3 (2.4-7.9) | |
| **Molecular subtypes (IHC)** | | | | | | |
| Luminal A | 17 | (20.7) | 109 | (44.0) | | |
| Luminal B/HER2 neg | 16 | (19.5) | 76 | (30.6) | | |
| Luminal B/HER2 pos | 12 | (14.9) | 16 | (6.5) | | <0.001 |
| HER2 positive | 17 | (20.7) | 12 | (4.8) | | |
| Triple negative | 20 | (24.4) | 35 | (14.1) | | |

n: number of patients; OR: odds ratio; CI: confidence interval; P-values by pearson's chi-square test. Missing data: histological type: n=3; tumor diameter: n=6; nodal status: n=10; ER: n=1; PR: n=1; HER2: n=23

**Supplementary table 2.** Age at diagnosis, clinico-pathologic data. METABRIC combined cohort (n=1784).

| | Age at diagnosis (years) | | | | | | |
|---|---|---|---|---|---|---|---|
| | <40 | | 40-49 | | ≥50 | | |
| | n (%) | | n (%) | | n (%) | | P |
| **Histological grade** | | | | | | | |
| 1 | 1 | (0.9) | 25 | (9.8) | 120 | (8.9) | <0.001 |
| 2 | 20 | (18.7) | 85 | (33.2) | 568 | (42.1) | |
| 3 | 86 | (80.4) | 146 | (57.0) | 660 | (49.0) | |
| **Tumor diameter** | | | | | | | |
| ≤2.0 cm | 33 | (31.4) | 91 | (35.4) | 414 | (29.4) | 0.16 |
| >2.0 cm | 72 | (68.6) | 166 | (64.6) | 992 | (70.6) | |
| **Nodal status** | | | | | | | |
| Negative | 36 | (33.6) | 136 | (52.1) | 750 | (53.0) | 0.001 |
| Positive | 71 | (66.4) | 125 | (47.9) | 666 | (47.0) | |
| **ER (IHC)** | | | | | | | |
| Positive | 35 | (33.0) | 158 | (61.2) | 1149 | (83.0) | <0.001 |
| Negative | 71 | (67.0) | 100 | (38.8) | 235 | (17.0) | |
| **Molecular subtype (PAM50)** | | | | | | | |
| Luminal A | 20 | (18.7) | 108 | (41.4) | 593 | (41.9) | <0.001 |
| Luminal B | 11 | (10.3) | 43 | (16.5) | 438 | (30.9) | |
| HER2 type | 21 | (19.6) | 34 | (13.0) | 185 | (13.1) | |
| Basal-like | 55 | (51.4) | 76 | (29.1) | 200 | (14.1) | |

n: number of patients; P-values by Pearson's chi-square test. Missing data: Histologic type: n=2; histologic grade: n=73; tumor diameter: n=16; ER: n=36

**Supplementary table 3.** Age at diagnosis and prognostic value of Ki67 TMA scoring. Bergen combined cohort (n=774).

| Ki67 | <40 (n=78) | | 40-49 (n=240) | | 50-59 (n=212) | | >60 (n=244) | |
|---|---|---|---|---|---|---|---|---|
| | Hazard ratio (CI 95%) | P value | Hazard ratio (CI 95%) | P value | Hazard ratio (CI 95%) | P value | Hazard ratio (CI 95%) | P value |
| **20 percentile (2.40 %)** | 0.84 (0.28 - 2.46) | 0.745 | 1.49 (0.71 - 3.12) | 0.295 | 2.64 (0.93 - 7.48) | 0.069 | 1.65 (0.70 - 3.89) | 0.253 |
| **30 percentile (4.05 %)** | 1.45 (0.42 - 4.97) | 0.558 | 1.98 (0.97 - 4.02) | 0.060 | 2.62 (1.09 - 6.34) | 0.032 | 2.34 (1.13 - 4.84) | 0.022 |
| **40 percentile (5.89 %)** | 1.16 (0.42 - 3.21) | 0.781 | 2.51 (1.33 - 4.74) | 0.004 | 2.15 (1.03 - 4.49) | 0.042 | 2.36 (1.22 - 4.54) | 0.011 |
| **50 percentile (8.00 %)** | 1.46 (0.52 - 4.05) | 0.471 | 2.64 (1.49 - 4.69) | 0.001 | 2.95 (1.44 - 6.06) | 0.003 | 1.87 (1.05 - 3.33) | 0.034 |
| **60 percentile (10.33 %)** | 1.51 (0.57 - 3.98) | 0.405 | 2.87 (1.66 - 4.95) | <0.001 | 3.27 (1.64 - 6.54) | 0.001 | 1.93 (1.08 - 3.45) | 0.026 |
| **70 percentile (15.39 %)** | 1.00 (0.41 - 2.47) | 0.993 | 3.26 (1.92 - 5.53) | <0.001 | 3.79 (1.93 - 7.44) | <0.001 | 1.50 (0.79 - 2.85) | 0.212 |
| **80 percentile (23.4 %)** | 0.94 (0.40 - 2.23) | 0.896 | 2.51 (1.51 - 4.15) | <0.001 | 3.80 (1.81 - 7.97) | <0.001 | 2.50 (1.30 - 4.83) | 0.006 |
| **Continuous** | 1.00 (0.98 - 1.01) | 0.745 | 1.02 (1.01 - 1.03) | <0.001 | 1.03 (1.01 - 1.04) | <0.001 | 1.02 (1.00 - 1.03) | 0.017 |

Ki67 scoring dichotomized into high and low using different cut-points. The percentiles represent all TMA-based scoring from the Bergen combined cohort. HR and p-values by univariate survival analyses by Cox' proportional hazards regression model.
